# Evolutionary quantitative proteomics of reproductive interactions in *Drosophila*

**DOI:** 10.1101/2022.02.04.479202

**Authors:** Martin D. Garlovsky, Yasir H. Ahmed-Braimah

## Abstract

Reproductive traits evolve rapidly between species. Understanding the causes and consequences of this rapid divergence requires characterization of male and female reproductive proteins and how their interaction mediates fertilisation success. Species in the *Drosophila virilis* clade exhibit interspecific postmating prezygotic reproductive isolation, making them ideal for studies on diversification of reproductive proteins and their role in speciation. Here we present a novel method to simultaneously identify and quantify transferred male ejaculate proteins and the female reproductive proteome using multiplexed Tandem-Mass-Tag (TMT) isobaric labelling of the lower female reproductive tract before and immediately after mating in three species of the *virilis* group. We identified over 200 putative male ejaculate proteins, many of which show differential abundance between species. We also identified over 2000 proteins that provide the first description of the female reproductive tract proteome outside of the *D. melanogaster* group. The female reproductive tract proteome shows significant divergence between species, especially female-specific serine-type endopeptidases, which showed differential abundance between species and elevated rates of molecular evolution that is similar to that of male seminal fluid proteins. We also found that only ~1/3 of male ejaculate proteins are likely produced in the accessory glands and code for a signal peptide sequence. Finally, we assessed the utility of species-specific compared to single species query databases for protein identification and quantification from multiple species’ samples. Our results show that using species-specific query databases dramatically improves protein identification and quantification.

## Introduction

The complex molecular interactions between male and female gametes and their accompanying reproductive fluid are critical for fertilisation success (1–5). Additional complexity arises in internally fertilising taxa compared to external fertilisers because fertilising sperm must traverse the often intricate labyrinth of the female reproductive tract. Moreover, frequent polyandry and extended periods of sperm storage presents the opportunity for postcopulatory sexual selection (6–8) and the need to maintain sperm viability and retention in storage (9). The reproductive fluid in the female reproductive tract and the male ejaculate consist of a complex mixture of carbohydrates, lipids, microbes and proteins (10–12). However, we still lack a comprehensive understanding of how the molecular composition of the reproductive fluid influences fertilisation success.

Proteins in the male and female reproductive fluid (i.e., the *reproductive proteome*) have important impacts for reproduction. For instance, in *Drosophila melanogaster*, over 200 male seminal fluid proteins (SFPs) have been identified, some of which are important in eliciting the postmating female response that affects female postmating physiology and behaviour (13, 14). Similarly, the fluid surrounding ova and other female reproductive tract secretions are critical in determining fertilisation success (3, 15–19). Yet, few studies have investigated the molecular composition and function of the female reproductive tract proteome despite requisite interactions between female and male molecules (16, 20–22).

Cooperative interactions between the sexes at the molecular level are essential for reproduction (13). For instance, the stereotyped postmating female response brought about by receipt of the male SFP Sex Peptide requires interaction with the female derived Sex Peptide Receptor in *D. melanogaster* (23,24). However, conflicts over reproduction that arise from the different reproductive interests of the sexes can lead to the evolution of male and female strategies to control reproduction (25, 26). For instance, females secrete proteases to digest the male ejaculate to maintain control over reproduction (27–29). In turn, the male ejaculate may contain protease inhibitors to counteract such female control (30). Thus, rapid molecular co-evolution of reproductive proteins can lead to species- or population-specific ejaculate × female interactions (1, 31, 32).

Co-diversification within populations may result in mismatched ejaculate × female reproductive tract interactions between populations, and the emergence of postmating prezygotic isolation (32, 33). Heterospecific ejaculates may fair worse in postcopulatory sexual selection (34, 35), have reduced motility (15, 36), or elicit an abnormal postmating female response, which can cause impaired sperm use or reduced oviposition rates (16, 20, 37, 38). A growing body of literature has begun characterising the genomic, transcriptomic, and proteomic differences between taxa that may contribute to postmating prezygotic isolation (16,20, 33, 39–43).

The rapid molecular evolution of reproductive proteins poses a significant challenge for protein identification and quantitation with high throughput proteomics using liquid-chromatography tandem mass-spectrometry (LC-MS/MS) and data dependent acquisition (DDA). Most proteomics studies use a single query database to assign mass spectra to peptides and proteins, either using the proteome of one of the species of interest (40), the proteome of the closest available relative (39), or performing *in silico* predictions to generate a reference proteome (44). Using a single species query database for cross-species comparisons may suffer from problems akin to read mapping bias in short read RNA sequencing (RNA-Seq) analysis pipelines (45). For instance, when using a protein database from another species, if a protein has undergone significant amino acid substitutions, spectra may be matched to an incorrect peptide or be unassigned completely, resulting in misidentification or inaccurate protein abundance estimation (46). Studies have revealed deep coverage of the sperm proteome across distantly related taxa after relatively long evolutionary time periods (46, 47). While the sperm proteome is likely subject to evolutionary constraint (48, 49), cross-species identification using a single species query database may be more challenging for reproductive proteins which evolve more quickly, such as SFPs.

Species in the *Drosophila virilis* clade (Fig. 1a) provide an ideal system in which to characterise divergence in the male ejaculate and female reproductive tract proteome. Postmating prezygotic isolation is pervasive in heterospecific crosses between these species, where sperm are rapidly lost from the female sperm storage organs (seminal receptacle and spermathecae) or become incapacitated, resulting in dramatically reduced fertilisation success (37, 50, 51). Previous studies have shown that SFPs evolve rapidly, species show divergent gene expression patterns in the male reproductive tract (42), and several rapidly evolving SFPs lie within paternal quantitative trait loci (QTL) that are associated with postmating prezygotic isolation (42). Furthermore, gene expression in the female reproductive tract is significantly perturbed after heterospecific mating (20).

**Fig. 1.**
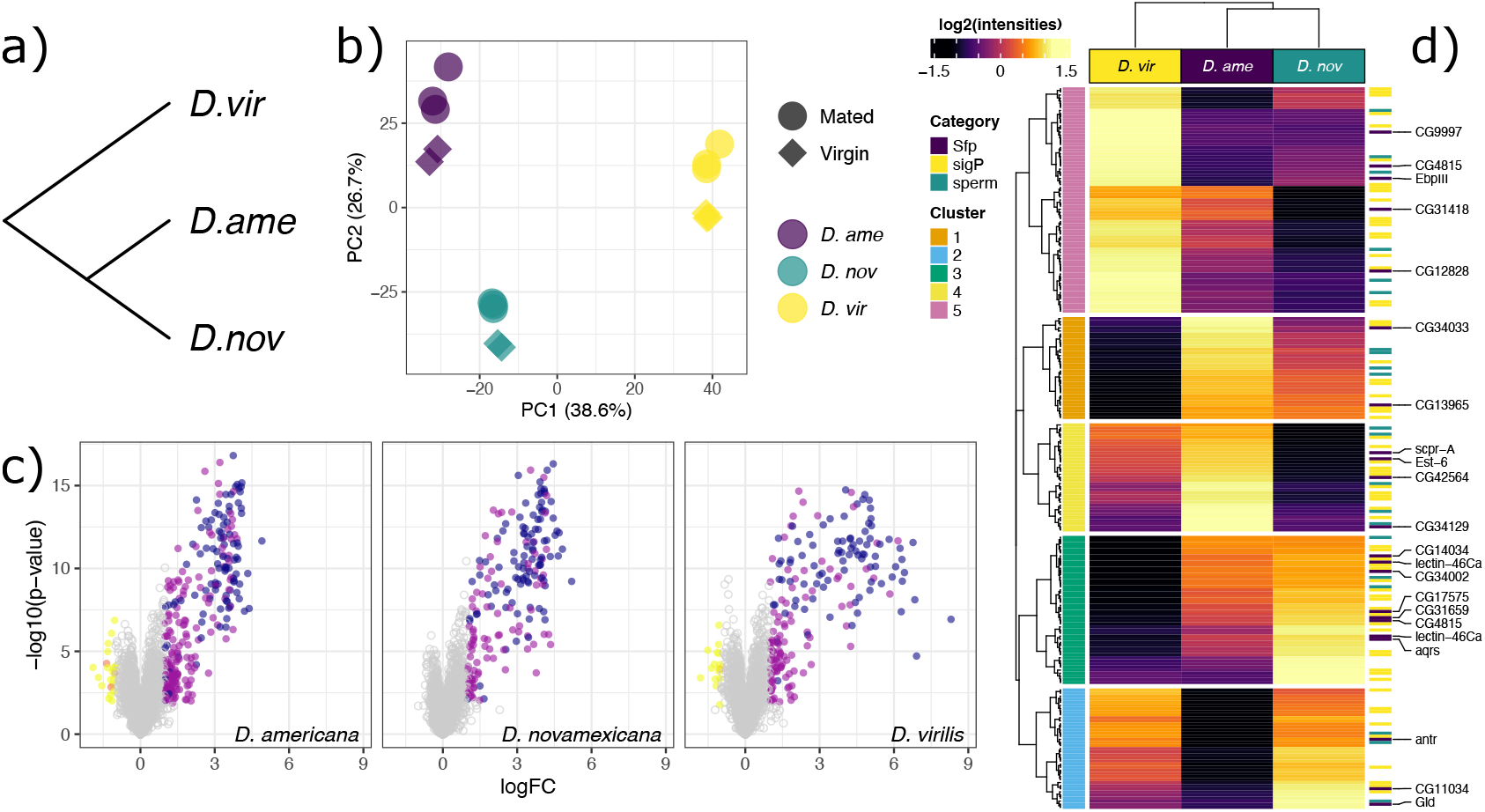
a) Phylogenetic relationship between species in the *virilis* group used in this study. b) Principal component analysis (PCA) plot showing sample relationships. PC1 explained 38.6% of the variance. PC2 explained 26.7% of the variance. c) Volcano plots demonstrating identification of “ejaculate candidate” proteins in each species. Positive values indicate higher abundance in mated samples. Coloured points show proteins higher abundance (log2-fold-change > 1 and FDR corrected p-value < 0.05) in mated samples with (blue) or without (purple) a signal peptide, or virgin samples with (orange) or without (yellow) a signal peptide. d) Abundance of ejaculate candidates (median centered log2(normalised abundances) averaged across replicates). K-means clusters are shown on the left. Signal peptides and *D. melanogaster* sperm and SFP orthologs are highlighted on the right. *D. melanogaster* SFP ortholog names are highlighted on the right. b) and d) use the combined species database (n = 2704), whereas c) uses the species-specific databases.

To identify male ejaculate proteins that are transferred during mating requires distinguishing proteins which are transferred to the female reproductive tract from other so called “house keeping” proteins found in the reproductive tract tissues. One successful approach is to use heavy isotopic labelling of one sex and distinguish mass spectra by a mass:charge shift. For instance, stable isotope labelling by amino acids in cell culture (SILAC) and similar techniques provided the first comprehensive list of transferred male seminal fluid proteins in *Drosophila* (52, 53). While SILAC has been the gold standard for identifying protein origin in a mixture of ≥2 sample, this method is time consuming, labour intensive and requires expensive growth media that may not be optimised for different species. Other approaches have recently been applied, e.g., Sepil et al. (54) identified over 50 previously unidentified male *D. melanogaster* ejaculate proteins by comparing the abundance of proteins in the accessory glands and ejaculatory ducts—the main source of male SFPs in insects (55)—of males before and after mating (54).

Here, we present a simple quantitative method to simultaneously characterise the transferred male ejaculate proteome and female reproductive tract proteome while avoiding the need for expensive and time consuming heavy labelling techniques. We compared protein abundances in the lower reproductive tracts of unmated females to those that were collected immediately after mating (thus containing ejaculate proteins) using tandem mass tag (TMT) labelled samples. Copulation duration in the *virilis* group is ~2 minutes and the majority of postmating female responses in *Drosophila* take place ~3 hours after the start of mating (20, 56), such that the female reproductive tract should remain in a “virgin-like” state.

Our objective is three-fold. First, to characterise the male ejaculate and female reproductive tract proteome of the *virilis* group, which is an emerging genetic model for understanding evolutionary mechanisms of reproductive interactions and speciation. Second, to identify differences between species in protein abundance in both the male ejaculate and the female reproductive tract to characterize the patterns of divergence in protein abundance. Finally, to assess the utility of different species query databases for protein identification and quantitation, where we predict greater precision using species-specific sequence information.

## Methods

### A. Fly stocks and maintenance

The following isofemale lines were used for each species: *D. americana* SB.02.06 (obtained from Bryant F. McAllister, University of Iowa); *D. novamexicana* 15010-1031.04, and *D. virilis* 15010-1051.49 (National Drosophila Species Stock Center, Cornell University). Flies were raised on cornmeal/sucrose media supplemented with yeast and kept at 22°C on a natural light cycle. We collected virgins within 2 days of eclosion and kept flies in single sex groups of 10-25 until sexually mature (~ 10 days old).

### B. Tissue collection and protein extraction

We paired virgin flies of the same species individually in food vials until copulation—which lasts ~2 minutes—was observed. We flash froze flies in liquid nitrogen within 30 seconds after copulation terminated and stored flies at −80°C until dissections. We thawed females and dissected the lower reproductive tract (uterus, seminal receptacle and spermathecae) in a drop of phosphate buffered saline (PBS) using fine forceps. We pooled 40-50 female reproductive tracts per replicate in 50μl lysis buffer (5% SDC; 1% SDS in HPLC grade water) kept on ice, and then freeze thawed pooled samples (×3) by placing on dry ice for 5 minutes, thawing to room temperature and then vortexing for 30 seconds. Finally, we centrifuged samples at 17,000G for 5 minutes at 4°C and collected the resulting supernatant. Samples were stored at - 80°C and shipped to the Cambridge Centre for Proteomics (Cambridge, UK) on dry ice for further processing.

### C. Liquid-chromatography tandem mass spectrometry

Liquid-chromatography tandem mass spectrometry (LC-MS/MS) was performed at the Cambridge Centre for Proteomics using ThermoFisher Scientific TMTpro 16plex Isobaric Label Reagent Set 0.5mg. Labelling was performed according to manufacturer instructions. 80μg of proteins per sample was labelled and protein estimation was done using RC-DC protein assay from Bio-Rad.

Labelled samples were cleaned on SepPack C18 cartridge from Waters before being fractionated on ACQUITY UPLC system using ACQUITY UPLC BEH C18 1.7um, 2.1 × 150mm column. Parameters of the chromatography method are as follows: flow 0.244ml/min, linear gradient starting from 95% of buffer A: 20mM ammonium formate pH10, 5% buffer B: 20mM ammonium formate in 80% acetonitrile pH10 ending at 100% of buffer B during the time of 75 minutes. PDA detector lamp setting: 210nm-400nm 1 minute fractions were collected starting from peptide elution observed on chromatogram. Fractions were concatenated: 1st fraction with the middle run fraction and so on to achieve a different elution profile per combined fraction later on when performing LC MS/MS. In total 15 fractions were produced for LC-MS/MS. Dried fractions from the high pH reverse-phase separations were resuspended in 30 μL of 0.1% formic acid and placed into a glass vial. 1 μL of each fraction was injected by the HPLC autosampler and separated by the LC method detailed below. Fractions were combined into pairs (i.e. the first fraction with the middle fraction etc.) and were analysed by LC-MS/MS.

LC-MS/MS experiments were performed using a Dionex Ultimate 3000 RSLC nanoUPLC (Thermo Fisher Scientific Inc, Waltham, MA, USA) system and a Lumos Orbitrap mass spectrometer (Thermo Fisher Scientific Inc, Waltham, MA, USA). Peptides were loaded onto a precolumn (Thermo Scientific PepMap 100 C18, 5mm particle size, 100Å pore size, 300 mm i.d. × 5mm length) from the Ultimate 3000 auto-sampler with 0.1% formic acid for 3 minutes at a flow rate of 10 μL/min. After this period, the column valve was switched to allow elution of peptides from the pre-column onto the analytical column. Separation of peptides was performed by C18 reverse-phase chromatography at a flow rate of 300 nL/min using a Thermo Scientific reverse-phase nano Easy-spray column (Thermo Scientific PepMap C18, 2mm particle size, 100A pore size, 75 mm i.d. × 50cm length). Solvent A was water + 0.1% formic acid and solvent B was 80% acetonitrile, 20% water + 0.1% formic acid. The linear gradient employed was 2-40% B in 93 minutes. Total LC run time was 120 mins including a high organic wash step and column re-equilibration.

The eluted peptides from the C18 column LC eluant were sprayed into the mass spectrometer by means of an Easy-Spray source (Thermo Fisher Scientific Inc.). All *m/z* values of eluting peptide ions were measured in an Orbitrap mass analyzer, set at a resolution of 120,000 and were scanned between *m/z* 380-1500 Da. Data dependent MS/MS scans (Top Speed) were employed to automatically isolate and fragment precursor ions by collision-induced dissociation (Normalised Collision Energy (NCE): 35%) which were analysed in the linear ion trap. Singly charged ions and ions with unassigned charge states were excluded from being selected for MS/MS and a dynamic exclusion window of 70 seconds was employed. The top 10 most abundant fragment ions from each MS/MS event were then selected for a further stage of fragmentation by Synchronous Precursor Selection MS3 (1) in the HCD high energy collision cell using HCD (High energy Collisional Dissociation, NCE: 65%). The *m/z* values and relative abundances of each reporter ion and all fragments (mass range from 100-500 Da) in each MS3 step were measured in the Orbitrap analyser, which was set at a resolution of 60,000. This was performed in cycles of 10 MS3 events before the Lumos instrument reverted to scanning the *m/z* ratios of the intact peptide ions and the cycle continued.

### D. Protein identification and quantitation

We processed .RAW data files using Proteome Discoverer v2.4 (Thermo Fisher Scientific) and Mascot (Matrix Science) v2.6. We first generated a custom reference proteome database for each species. Briefly, we mapped available RNA-Seq data from male reproductive tissues of the three species to their respective genomes (42) and assembled their transcriptomes using StringTie (**?**). For *D. novamexicana* and *D. virilis*, we used the recently developed/updated NCBI RefSeq assembly and gene annotations (GCF_003285875.2 and GCF_003285735.1, respectively) as a guide in transcriptome assembly, and we used a recently generated PacBio genome of *D. americana* (https://github.com/YazBraimah/Dvirilis_group_PacBio_genomes). Assembled transcripts were annotated and protein coding genes were predicted with TransDecoder (https://github.com/TransDecoder). The total number of protein isoforms was as follows: *D. americana* (77,209 entries), *D. novamexicana* (140,262 entries), and *D. virilis* (183,293 entries). We searched raw files iteratively against the custom reference proteomes for each species and the common repository of contaminant proteins (cRAP, 125 sequences). We specified trypsin as enzyme with a maximum of 2 missed cleavage, 16-plex TMTpro and carbamidomethyl (C) as fixed modification and oxidation (M) and deamidation (NQ) as variable modifications. We used Percolator (57) to estimate the false discovery rate (FDR) and applied stringent thresholds for peptide (FDR < 0.01) and protein identification (FDR < 0.05). Protein identification allowed an MS tolerance of ±10 ppm and an MS/MS tolerance of ±0.8Da ppm along with permission of up to 2 missed tryptic cleavages. Quantification was performed in Proteome Discoverer on the 16 reporter ion intensities per peptide by calculating the sum of centroided reporter ions within a ±2 millimass unit (mmu) window. Mass spectrometry data are available via the ProteomeXchange Consortium (ID: XXXXXX).

### E. Combined database compilation

We identified 1:1 reciprocal orthologs between all three species using OrthoFinder (58, 59), with the three species complete proteome sequence files as input and default settings. We then combined species-specific abundance data for each protein by “orthoGroups”.

### F. Experimental design and statistical rationale

In total we collected 16 samples for labelling using the 16-plex TMT reagent kit. We collected three biological replicates for mated samples for each species, and *D. virilis* virgin samples, and two replicates each for *D. americana* and *D. novamexicana* virgin samples. We chose this design to increase replication of the virgin *D. virilis* samples, which we expect to have diverged most from the sister species *D. americana* and *D. novamexicana*. We also expect greater differences between mated samples due to the rapid evolution of male seminal fluid proteins.

We applied stringent thresholds for peptide (FDR < 0.01) and protein identification (FDR < 0.05) and ensured accuracy of quantitative estimates using strict mass tolerance and retention time cut-offs. We performed differential abundance analyses between samples using the edgeR package (60) in R v.4.0.3 (61) (see Statistical analysis). For differential abundance analyses we filtered proteins identified by two or more unique peptides and normalised abundance data using the trimmed mean of M-values (TMM) method. Differences in protein abundance were determined based on a log2 fold-change > |1| and FDR corrected *p*-value < 0.05. We inspected diagnostic plots for all analyses and corrected *p*-values for multiple testing using the Benjamini-Hochberg method (62).

### G. Statistical analysis

We performed all statistical analyses in R v.4.0.3 (61). Complete code and analysis can be found on GitHub. For differential abundance analyses, we filtered proteins identified by fewer than two unique peptides and used *voom* normalisation and fit protein-wise linear models (*lmFit*) with empirical Bayes smoothing (*eBayes*) using edgeR (60).

### H. Evolutionary rates analysis

We obtained previously published pairwise rates of non-synonymous (dN) to synonymous (dS) nucleotide substitutions (dN/dS) (20) and maximum-likelihood estimates of amino acid substitutions on the *virilis* subspecies phylogeny (*ω*) (42) that were calculated using PAML (63).

### I. Gene ontology (GO) analysis

We performed gene ontology enrichment analysis using the ClueGO plugin v.2.5.7 (64) for Cytoscape v.3.8.0 (65) and *D. virilis* FlyBase gene numbers (FBgns). We performed right-sided hypergeometric enrichment tests, and applied multiple testing correction using the Benjamini-Hochberg method. All GO terms reported had a *p*-value <0.01. Specific network settings can be found in figure and table legends.

## Results

We retrieved 2704 reciprocal orthologs between all three species for proteins identified by LC-MS/MS to compile the combined database (Table S1). This accounts for over 64% of proteins identified using each species query database, which is significantly lower than the percentage of all proteins in the proteome assigned to orthogroups using OrthoFinder (88%), potentially arising from the rapid evolution of reproductive genes (66). We report results from the combined database unless otherwise stated. Principal component analysis showed that the first three components explained > 83% of the variance. PC1 (38.6% of variance explained) separated species, with the largest difference between *D. virilis* and the sister species *D. americana* and *D. novamexicana* (Fig. 1a,b). PC2 and PC3 explained 26.7%, and 17.9% of the variance, respectively, further separating virgin from mated samples (Fig.1b).

### J. Male ejaculate proteome

We identified 206 proteins that were significantly more abundant in mated samples in at least one species using the combined database (henceforth “ejaculate candidates”; i.e., putative SFPs and sperm proteins), which account for 8% of all identified proteins (Fig. 1c). The majority of ejaculate candidates identified (92) were shared between all three species (55-74%), i.e., representing the *core ejaculate proteome* (Fig. S1a). Over half (66%) of ejaculate candidates had a signal peptide sequence, indicative of potentially secreted proteins (Fig. 1c,d). To compile a complete list of ejaculate candidates for further analysis, we combined all ejaculate candidates with orthologs identified using each species database (Table S2). Of these 228 proteins with orthologs, the majority (196; 86%) were identified as ejaculate candidates in at least two out of three databases (Fig. S1b). The remaining, database-specific ejaculate candidates, which account for between 29-40% of ejaculate candidates identified using a single species database (Fig. S1c), did not have orthologs precluding further comparisons.

Ejaculate candidates included orthologs of 21 *D. melanogaster* sperm proteins (49, 67) and 55 *D. melanogaster* SFPs (14), including 5 members of the sex peptide network; Antares (*CG30488*), Aquarius (*CG14061*), Lectin-46Ca (*CG1656*), CG9997, and CG17575 (68). Additionally, 52 proteins overlap with genes that were inferred to be SFPs by mRNA expression or otherwise showed accessory gland or ejaculatory bulb biased expression (Fig. S2; (42)). Ejaculate candidates were enriched for GO terms expected for sperm and seminal fluid proteins, including metabolic and catabolic processes, proteolysis, peptidase activity (BP), extracelluar space (CC), enzyme inhibitor activity, (serine-type/endo-)peptidase activity and hydrolase activity (MF) (Table S3).

It has been suggested that transferred male ejaculate proteins may be “swamped out” by the more abundant proteins in the female reproductive tract (54). The average abundance of ejaculate candidates was not significantly different from the remaining (female reproductive tract) proteome for *D. americana* (Mann-Whitney U test, *p* = 0.06), or *D. novamexicana* (p = 0.29) but was lower for *D. virilis* (p < 0.001) (Fig. S3).

#### J.1. Ejaculate divergence

To identify differences in the transferred male ejaculate proteome between species we performed differential abundance analyses between each species pair using all ejaculate candidates (n = 228). There were fewer differentially abundant proteins between the more lOQFC Lrļ —— CG11034 closely related *D. americana* and *D. novamexicana* (59; 28%) than between the more distantly related pairs *D. americana* and *D. virilis* (100; 44%) or *D. novamexicana* and *D. virilis* (97; 43%) (Table 1; Table S4). Proteins with predicted signal peptides were not more likely to be differentially abundant in all comparisons (Fisher’s exact tests, all *p* > 0.88), as were *D. melanogaster* sperm proteins (Fisher’s exact tests, all *p* > 0.489; Fig. 1d). Differentially abundant ejaculate candidates again showed GO enrichment (*p* < 0.01) for metabolic processes (BP); organelle subcompartment (CC); and acid phosphatase activity (MF). Using a less stringent *p*-value cutoff (*p* < 0.05) serine-type endopeptidase inhibitor activity and peptidase inhibitor/regulator activity were also enriched. Spermatid development (*p* = 0.059) was also notable among enriched terms, indicating that we identified some sperm proteins in our samples (Table S5).

**Table 1.** Summary of numbers of proteins identified using each species database and the combined database (CDB). Numbers of differentially abundant proteins between species for ejaculate candidates and virgin female reproductive tracts. For the combined database the number of proteins identified by two unique peptides was calculated using at least one search database.

|  | Database |  |  |  |
| --- | --- | --- | --- | --- |
|  | *D. ame* | *D. nov* | *D. vir* | CDB |
| Total no. proteins | 4223 | 4160 | 4051 | 2704 |
| Two unique peps. | 4020 | 3899 | 3785 | 2645 |
| Ejac. candidates | 346 | 322 | 264 | 206 |
| Ejac. candidates (%) | 8.6 | 8.3 | 7.0 | 7.8 |
| Ejaculate divergence |  |  |  |  |
| *D. ame* vs. *D. nov* | 57 | 53 | 34 | 59 |
| *D. ame* vs. *D. vir* | 155 | 140 | 101 | 100 |
| *D. nov* vs. *D. vir* | 177 | 177 | 110 | 97 |
| Female reproductive tract div. |  |  |  |  |
| *D. ame* vs. *D. nov* | 158 | 153 | 133 | 327 |
| *D. ame* vs. *D. vir* | 384 | 363 | 382 | 455 |
| *D. nov* vs. *D. vir* | 408 | 404 | 387 | 353 |

Using k-means clustering identified 5 clusters of proteins that distinguish species in terms of protein abundance. Cluster 1 comprised proteins more abundant in *D. americana* compared to *D. virilis*; cluster 2 proteins lower in abundance in *D. americana* compared to both *D. novamexicana* and *D. virilis*; cluster 3 proteins lower in abundance in *D. virilis* compared to both *D. americana* and *D. novamexicana*, and cluster 5 proteins more abundant in *D. virilis* compared to both *D. americana* and *D. novamexicana*; finally, cluster 4 differentiated between *D. americana* and *D. novamexicana* (Fig. 1d; Fig. S4). *D. novamexicana* contained the largest number of *D. melanogaster* SFP orthologs in this set (11) that show higher relative abundance than *D. americana* and *D. virilis*; the remaining 10 SFP orthologs were either higher in abundance in *D. americana* or *D. virilis*. These results show that SFP abundance—which is not often considered when analyzing SFP function—is transient between species and may confer functional consequences.

### K. Female reproductive tract proteome

The majority of proteins showed no difference in abundance between mated and virgin samples (Table S6), including over 90% (197/218) of *D. melanogaster* sperm proteins we identified. We identified 52 female reproductive tract biased genes and 106 proteins that were previously found to show significant changes in gene expression after mating (20) (Fig. 2), only 5 of which showed differences in abundance between virgin vs. mated samples. Only two proteins with serine-type endopeptidase annotation were more abundant in virgin samples (FBgn0211434 in *D. americana*, FBgn0205380 in *D. virilis*), suggesting the female reproductive tract proteome had not yet undergone significant postmating changes in our samples—as expected. Overall, female reproductive tract proteins showed GO enrichment for various metabolic processes, cellular transport and organisation, gamete generation and development (BP), organelle and membrane components (CC), hydrolase and enzyme activity, and many molecule binding terms (MF) (Table S7). Twenty-two proteins were more abundant in virgin compared to mated samples (Table S8). However, no GO terms reached statistical significance (*p* < 0.05) for these proteins.

**Fig. 2.**
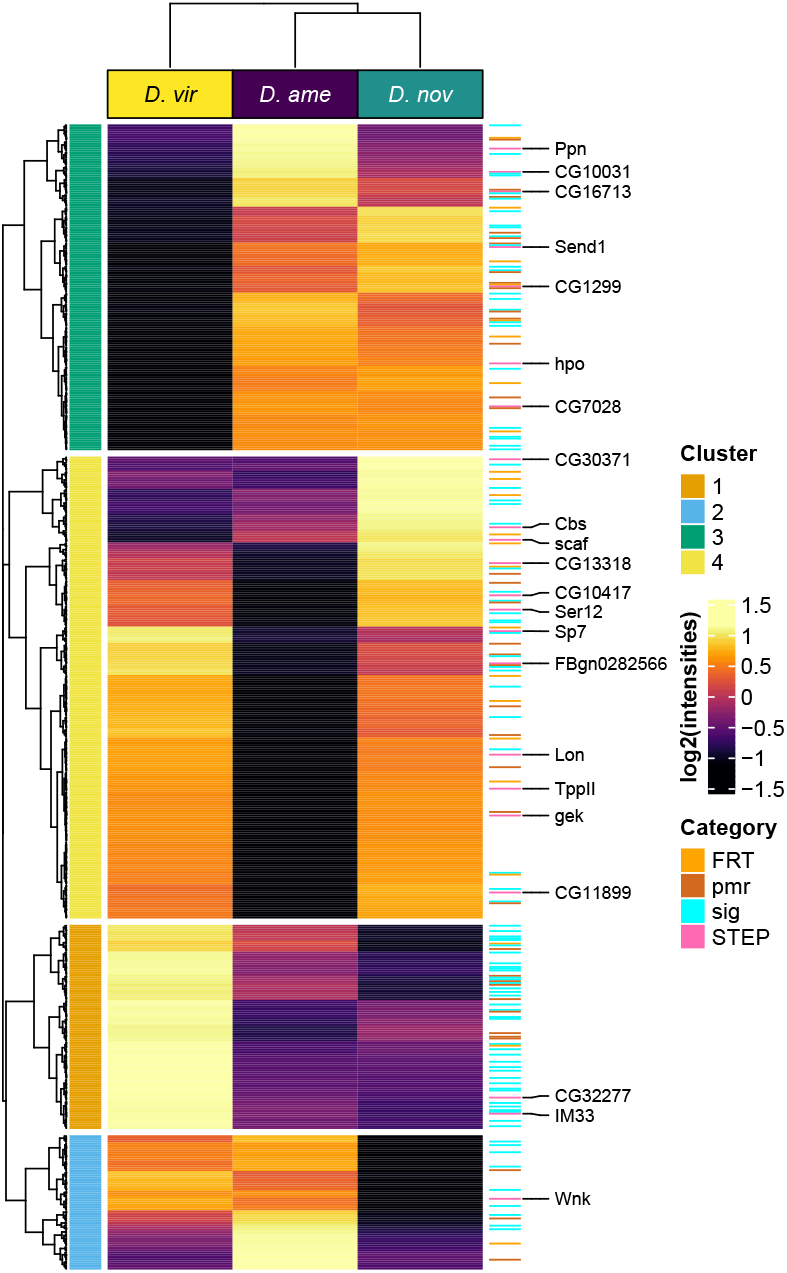
Differentially abundant female reproductive tract proteins between species (n = 629). K-means clusters (k = 4) are shown on the left. Female reproductive tract biased genes (FRT; brown), postmating response genes (pmr; orange), signal peptides (sig; blue), and serine-type endopeptidases (STEP; pink) are highlighted on the right. Values are median centered log2(normalised abundances).

We identified 323 signal peptides in the female reproductive tract proteome (Table S9), likely secreted into the female reproductive tract lumen and thus may directly interact with the male ejaculate. Secreted female reproductive tract proteins showed GO enrichment for extracellular space and matrix (CC) and hydrolase activity (MF) (Table S10). Serine-type (endo-)peptidase activity, regulation of proteolysis (BP), and serine-type endopeptidase activity, tetrapyrrole binding, hydrolase activity, and calcium ion binding (MF) showed enrichment at *p* < 0.05 (Table S10).

#### K.1. Female reproductive tract divergence

We performed differential abundance analysis between virgin samples after excluding ejaculate candidates (n = 2417; Table S6). Signal peptides were not more likely to be differentially abundant between *D. americana* and *D. novamexicana* (Fisher’s exact test, *p* = 0.20), but were more likely to be differentially abundant in comparisons involving *D. virilis* (both *p* < 0.001) (Fig. 2). Differentially abundant female reproductive tract proteins showed GO enrichment for many metabolic and catabolic processes, response to oxidative stress, chemical homeostasis, generation of precursor metabolites and energy (BP); actin cytoskeleton, endomembrane system, neuron projection, peroxisome, and cytoplasm (CC); many molecule binding terms, endopeptidase inhibitor activity, and (regulation of) endopeptidase activity (MF) (Table S11).

Using k-means clustering identified 4 clusters of proteins differentiating between species. Cluster 1 comprised proteins more abundant in *D. virilis* compared to both *D. americana* and *D. novamexicana*; cluster 2 proteins lower in abundance in *D. novamexicana* compared to both *D. americana* and *D. virilis*; cluster 3 proteins lower in abundance in *D. virilis* compared to both *D. americana* and *D. novamexicana*; and finally cluster 4 proteins lower in abundance in *D. americana* compared to both *D. novamexicana* and *D. virilis* (Fig. 2; Fig. S5).

### L. Evolutionary rates

Ejaculate candidates with a signal peptide sequence showed elevated dN/dS ratios compared to the remaining ejaculate candidates (Mann-Whitney U test, *p* < 0.001 in all comparisons) and the genome average (*p* < 0.006 in all comparisons), as expected for seminal fluid proteins (39, 42) (Fig. 3). Ejaculate proteins with a signal peptide and serine-type endopeptidase annotation evolved at a similar rate to other proteins with signal peptides (*p* > 0.29 in all comparisons). The remaining male ejaculate proteins were evolving more conservatively than the genome average (*p* < 0.028 in all comparisons; Fig. 3).

**Fig. 3.**
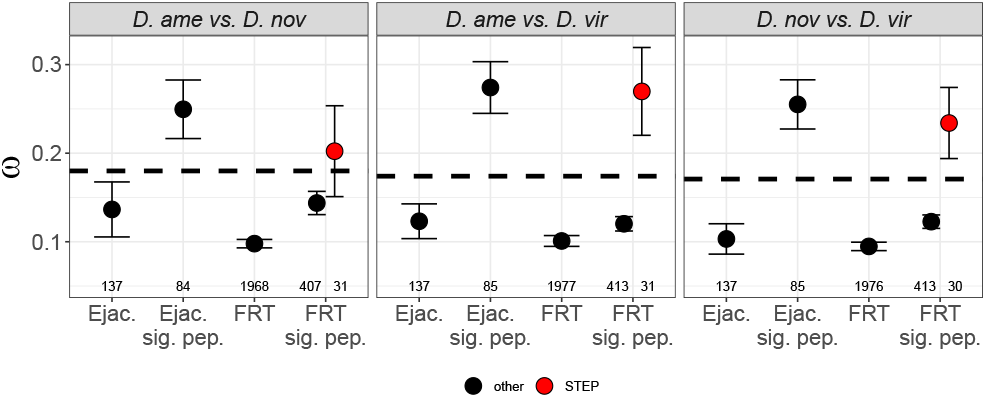
Pairwise non-synonymous to synonymous substitution rates (*ω*; mean ± standard error) for ejaculate candidates and female reproductive tract proteins with or without a signal peptide sequence. Female reproductive tract proteins with a signal peptide sequence and serine-type endopeptidase (STEP) annotation are shown in red. Numbers below points indicate sample sizes. Dashed line represents the genome average *ω* in each comparison (*D. americana* vs. *D. novamexicana* = [mean ± s.e.] 0.180 ± 0.003, n = 12687; *D. americana* vs. *D. virilis* = 0.174 ± 0.003, n = 12839; *D. novamexicana* vs. *D. virilis* = 0.171 ± 0.002, n = 12824).

Female reproductive tract proteins with a signal peptide sequence were evolving faster than the remaining female reproductive tract proteins (*p* < 0.001 in all comparisons), but more slowly than the genome average (*p* < 0.02 in all comparisons), except between *D. americana* and *D. novamexicana* (p = 0.073; Fig. 3). Female reproductive tract proteins with serine-type endopeptidase annotation and a signal peptide sequence showed elevated dN/dS compared to other secreted female reproductive tract proteins (*p* < 0.002 in all comparisons), and in all comparisons were evolving as fast as ejaculate proteins (*p* > 0.620 in all comparisons; Fig. 3). These secreted female reproductive tract proteins were also evolving faster than the genome average between *D. americana* and *D. virilis* (*p* = 0.003) and between *D. novamexicana* and *D. virilis* (*p* = 0.003), but not between *D. americana* and *D. novamexicana* (*p* = 0.489).

### M. Tissue-biased mRNA expression patterns of ejaculate proteins

Seminal fluid proteins are thought to be primarily produced in the accessory glands and ejaculatory duct and bulb and are thus expected to show tissue biased mRNA expression in these tissues. We examined the mRNA expression profile of ejaculate candidates in male reproductive tract tissues using mRNA expression data from Ahmed-Braimah et al. (42). First we classified ejaculate proteins based on their mRNA expression profile in the accessory glands, ejaculatory bulb, and testes; we considered the mRNA tissue-biased if it had higher expression in the focal tissue compared to all other reproductive male tissues and the gonadectomized male carcass. We identified 89, 87, and 78 genes that code for ejaculate proteins with a predicted signal peptide in *D. americana, D. novamexicana*, and *D. virilis*, respectively (Fig. 4A,D). Of those, 56, 26, and 22 show biased expression in the accessory glands of their respective species. Only a handful showed expression bias in the ejaculatory bulb (6, 3, and 3, respectively; Fig. 4B,D), and roughly a similar number to those that are tissue-biased in the accessory glands were testes-biased, albeit most of these did not contain a signal peptide (Fig. 4C,D). Importantly, ejaculate proteins with predicted signal peptides vastly outnumber non-secretory ejaculate proteins among the accessory gland-biased genes (3-7 fold; Fig. 4A). Surprisingly, however, the majority of ejaculate proteins (~2/3) show ubiquitous mRNA expression in all of the profiled male tissues, making it difficult to assess the exact source of a large number of proteins. These results suggest that a core set of ejaculate proteins—those with a secretion signal and accessory gland biased mRNA expression— are likely the main protagonists in reproductive interactions with the female reproductive tract.

**Fig. 4.**
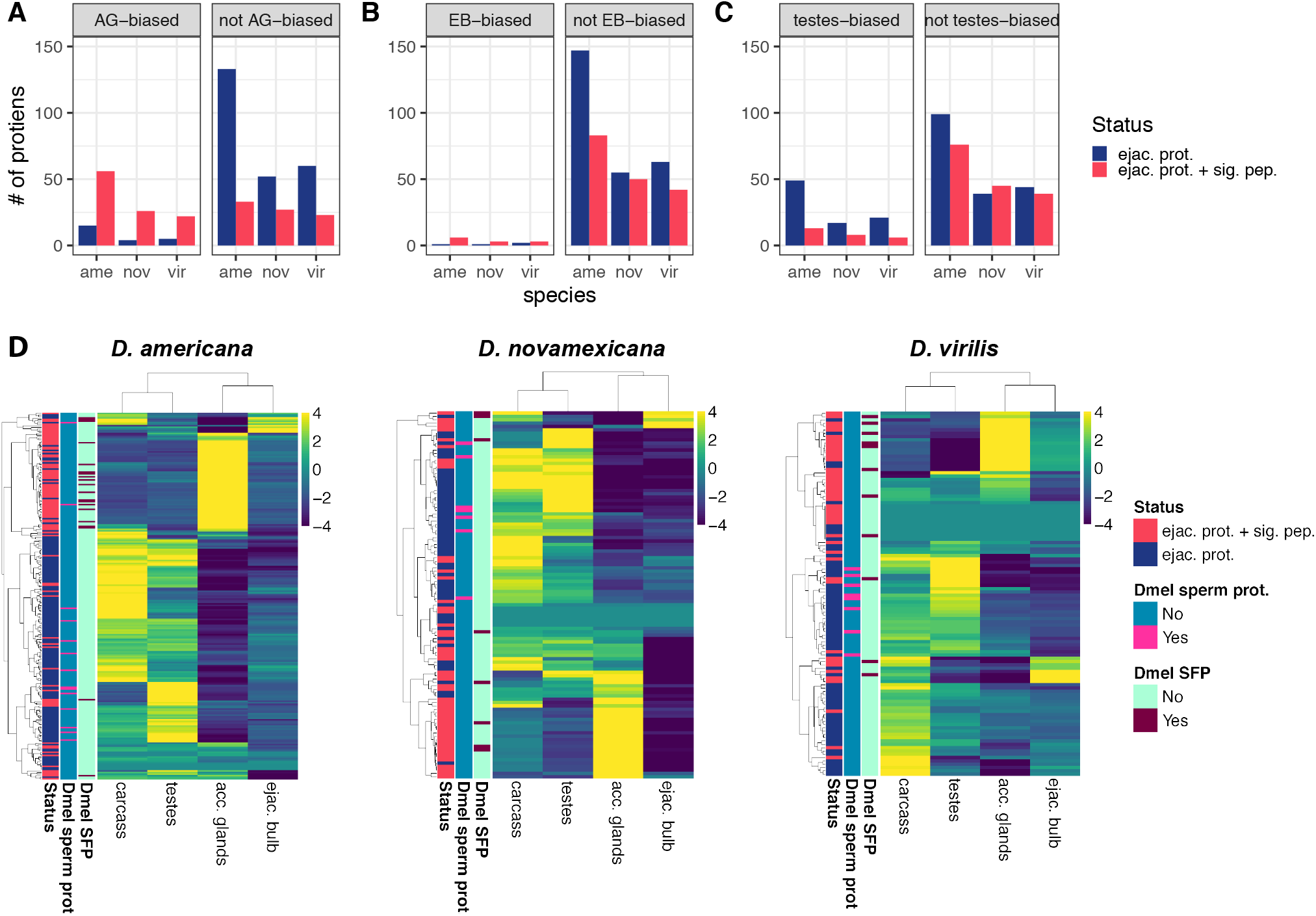
Dynamics of mRNA expression of transcripts that code for ejaculate proteins. (A-C) Number of ejaculate proteins classified by their maximum mRNA expression in the accessory glands (AG), ejaculatory bulb (EB), and testes for each species. (D) mRNA expression heatmaps of ejaculate proteins across the four male tissues for each of the three species. Annotation bars on the left indicate secretion signal status of each protein and whether the protein is orthologous to a *D. melanogaster* sperm protein or SFP (see legend).

Previous work that examined the correlation between mRNA and protein abundance has often found scant correlations between the two (69, 70). Here we explored the relationship between mRNA abundance in male tissues and protein abundance of transferred ejaculate proteins and find varying correlations depending on the tissue examined (Fig. S6). The highest Pearson correlations were found between mRNA transcripts that are expressed in the accessory glands or ejaculatory bulb and ejaculate proteins with a secretion signal (*R*=0.22-0.49 in all species), but the correlations were only significant in two of the three species for each tissue (Fig. S6). These results indicate that while the correlations between mRNA and protein abundance are weak, some tissues show significantly higher correlations than others and that these tissues likely represent the source of transferred ejaculate proteins.

### N. Database comparisons

In total we identified 4223 (Table S12), 4160 (Table S13), and 4051 (Table S14) proteins using the *D. americana, D. novamexicana*, and *D. virilis* query databases, respectively, with more than 93% of proteins identified by two or more unique peptides (Table 1). To assess variation in protein abundance between biological replicates we calculated the coefficient of variation (CV; 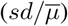) for each treatment (mated or virgin within each species). Mean CVs for each treatment were similar using each database (all < 0.032; Fig. S7), reflecting the high precision attained using TMT labelling. To assess variation attributable to differences in quantitation between databases we then calculated CVs for each replicate across all three species databases. Mean CVs were higher than compared to within each database (0.047-0.049), indicating more variability in quantitation between each species database for the same replicate than between biological replicates using a single database (Fig. S8).

To compliment this analysis, we compared the correlations in protein abundances between biological replicates of each treatment. If alternate species query databases are suitable for estimating protein abundances, then abundance correlations between the same replicate using alternate species query databases should theoretically approach unity. Conversely, if mass-spectra are not assigned accurately when using an alternative species database, then abundance estimates will be less correlated between databases. Mean Pearson’s correlations for each treatment were all > 0.98 when using a single species query database, indicating biological replicates behaved consistently (Fig. 5a,b). However, mean correlations ranged from between 0.79-0.88 when comparing protein abundances using alternate species databases to those obtained using species-specific query databases (Fig. 5a,b), suggesting alternate species databases provide less accurate protein abundance estimates. The greatest disparity was found when comparing abundance values for *D. americana* using the *D. virilis* query database and *vice versa* (Fig. 5b).

**Fig. 5.**
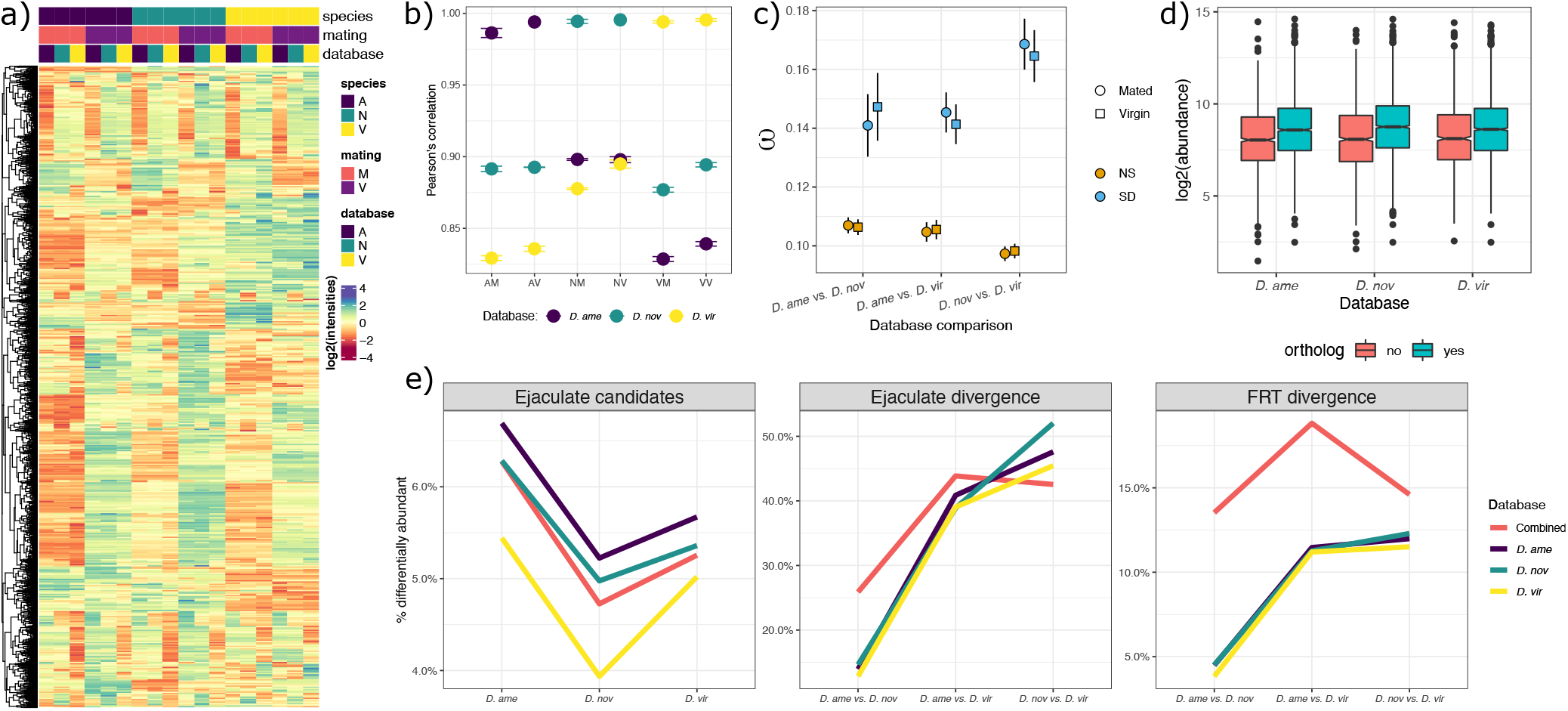
Database comparisons. a) Heatmap of average abundances for each treatment (mated and virgin) for each species using each species query database (n = 2699). b) Pearson’s correlations (means ± standard error) comparing protein abundances quantified using each species query database. Correlations were calculated between replicates within each treatment for each species for comparison’s within species, whereas correlations were calculated only between the same replicate when using alternate species query database. c) Mean dN/dS (± s.e.) comparing proteins showing differential abundance (SD; blue) between databases or not (NS; orange). d) Abundance of proteins identified using each database with- or without-reciprocal 1:1:1 orthologs. e) Proportion of proteins identified as ejaculate candidates (left); differentially abundant ejaculate candidates between species (middle); and female reproductive tract proteins between species (right). Proportions were calculated using the total number of differentially abundant proteins identified in each class divided by the total number of proteins used in each analysis (see Table 1). Note difference in Y-axis scales in (e). First letter: A, *D. americana*; N, *D. novamexicana*; V, *D. virilis*. Second letter: M, mated; V, virgin.

We then tested for differences in abundance when using alternative species databases which revealed between 816% of proteins differed in estimated abundance. Proteins detected at significantly different abundances had elevated dN/dS (Fig. 5c), as expected if amino acid substitutions reduce accuracy in peptide and protein assignment. Furthermore, proteins without reciprocal orthologs were found at lower average abundance (Fig. 5d), indicating less abundant proteins may be missed using an alternative species database.

We identified similar proportions of proteins that are more abundant in mated vs. virgin samples using all databases (Table 1). For ejaculate candidates, the *D. virilis* database consistently identified the lowest proportion of differentially abundant proteins between species, whereas the combined database identified more differentially abundant proteins between *D. americana* and *D. novamexicana*. For female reproductive tract proteins, the combined database identified considerably more differentially abundant proteins in all comparisons, while each species database identified similar proportions of differentially abundant proteins (Fig. 5e).

These analyses reveal the extent of bias that can be introduced when using divergent protein databases (i.e., interspecies databases) when querying mass spectra, and suggests that a species-specific query database is the best option for nearly complete and accurate protein identification and quantitation.

## Discussion

To understand the molecular interactions that mediate fertilisation success and postmating prezygotic isolation requires identifying differentiation in the reproductive proteome of both mating partners. Using a novel approach, we simultaneously characterised the male ejaculate and female reproductive tract proteome for three species in the *virilis* group. We identified a number of differentially abundant proteins between species, including endopeptidases and their inhibitors; male and female derived proteins which must interact to ensure normal fertility. Female serine-type endopeptidases were evolving as fast as male seminal fluid proteins, implicating rapid diversification of this class of proteins in the emergence of postmating prezygotic isolation. Finally, we assessed the utility of different species query databases for protein identification and quantitation. Our analysis shows that species-specific proteome data provides a more sensitive approach for detecting differentially abundant proteins. We caution against using a single species reference proteome for comparative evolutionary proteomics using data-dependent acquisition.

Identifying transferred male ejaculate proteins has long presented a challenge for research the investigates seminal fluid proteins. By comparing mated vs. virgin samples our method allowed us to identify a large number of putative SFPs without the need for expensive and labour intensive labelling techniques. That over half of all proteins that are more highly abundant in mated samples were secretory molecules suggests that our approach is effective at detecting seminal fluid proteins.

Some “postmating” female responses may be induced by courtship or during mating. Therefore, some of the differentially abundant proteins between mated and virgin samples may have been contributed by the female rather than transferred by the male. However, few proteins previously identified as characteristic of the postmating female response or showing female reproductive tract bias (20) showed differences in abundance, consistent with the finding that the majority of postmating female responses in *Drosophila* take place ~3 hours postmating (20). Further studies are necessary to establish the molecular changes involved in any “pericopulatory” female response.

We identified over 200 candidate ejaculate proteins (seminal fluid proteins and sperm proteins), including many previously identified putative SFPs from the *virilis* group, and orthologs of D. *melanogaster* SFPs and sperm proteins. Several orthologs of *D. melanogaster* SFPs that show differential abundance between species are notable for their effects on sperm entry and exit from storage (Fig. 1b). Ejaculatory bulb protein III (FBgn0011695) is integral to formation of the posterior mating plug which affects female sperm storage and fertility (71). Glucose dehydrogenase (FBgn0001112) affects sperm uptake and release from the spermathecae (72). Lectin-46Ca (FBgn0040093) affects long term sperm release from storage (73). Although we did not detect Sex Peptide, we did identify several other members of the Sex Peptide network (including Lectin-46Ca). Sex Peptide induces the long term postmating female response in *D. melanogaster*, and requires interactions among various members of the Sex Peptide network for binding of Sex Peptide to sperm (68). Together with the knowledge of rapid loss of sperm from storage as the mechanism resulting in reduced fertilisation success (37), our findings suggest that males transfer species-specific proportions of seminal fluid proteins during copulation. As such, females may not process a heterospecific ejaculate effectively, or receipt of an abnormal cocktail of seminal fluid proteins may elicit an abnormal postmating female response (20). Proteins that are involved in spermatid development were also among the differentially abundant ejaculate candidates.

The molecular composition of the female reproductive tract has historically not received the same level of interrogation as the male ejaculate. Our analysis expands on a growing effort to characterise the female reproductive tract proteome (16, 19, 28, 56). Interestingly, the majority (> 90%) of *D. melanogaster* sperm protein orthologs we identified showed no difference in abundance between mated vs. virgin samples, suggesting expression in the female reproductive tract. We identified over 100 female reproductive tract proteins with a signal peptide sequence. These secreted proteins likely interact with, and process, the male ejaculate in the female reproductive tract lumen. Accordingly, secreted female reproductive tract proteins were enriched for processes and functions relating to molecule binding, proteolysis and enzyme activity. Prominent among them were proteins with peptidase activity, which act to cleave other proteins (see below) (74–76). Secreted female reproductive tract proteins were also more likely to be differentially abundant between species in comparisons involving *D. virilis*, suggesting female proteins directly involved in ejaculate × female reproductive tract interactions can potentially underlie postmating prezygotic isolation.

Male seminal fluid proteins are predicted to play a disproportionate role in postmating prezygotic isolation due to their rapid diversification. This is evident in the much greater proportion of differentially abundant ejaculate candidates between species (> 25%), compared to the more modest numbers of proteins differing in abundance between female reproductive tract proteomes (< 25%), despite there being an order of magnitude more proteins for comparison. Unsurprisingly, however, we did identify a host of differentially abundant female reproductive tract proteins with orthologs linked to fertility in *D. melanogaster*. For instance, Send1 (FBgn0031406) is expressed in the spermathecae and recruits sperm into storage and reduces sperm motility in the seminal receptacle (77). Odorant binding proteins such as Odorant binding protein 56d (FBgn0034470) may play a role in sperm activation in insects (78). We also found enrichment of differentially abundant proteins that are involved in oxidative stress and metabolic processes. For instance, Hexokinase A (FBgn0001186) is involved in glucose homeostasis; Superoxide dismutase 3 (FBgn0033631) protects against reactive oxygen radicals; and Immune-regulated catalase (FBgn0038465) is involved in oxidant detoxification. Differences in the redox environment of the female reproductive tract suggests a potential mechanism that impedes sperm survival or motility within a heterospecific female reproductive tract (79). Finally, Immune induced molecule 33 (FBgn0031561) is secreted in to the haemolympth upon microbial infection. The potential role of microbes transferred during mating to postmating prezygotic isolation (12), or nevertheless that females mount an immune response against a heterospecific ejaculate requires further investigation.

Female reproductive tract genes evolve more conservatively than male reproductive genes and the genome average in the *virilis* group (20). However, we found that female serine-type endopeptidases evolved at an elevated rate, similar to that of male seminal fluid proteins. Serine-type endopeptidases mediate the cleavage of seminal fluid proteins in the female reproductive tract that are required to properly stimulate the postmating female response (74–76). We also found serine-type endopeptidases and their inhibitors were prominent among differentially abundant female reproductive tract proteins, and ejaculate candidates, respectively. Sequence and abundance divergence between species of endopeptidases and their inhibitors has been implicated in postmating prezygotic isolation across a range of taxa (16, 27, 28, 39, 80). Further analyses using divergence and polymorphism data are required to determine the evolutionary forces contributing to the fast molecular evolution of male and female derived proteins that contribute to postmating prezygotic isolation (81–84).

Our analysis of mRNA expression patterns among ejaculate proteins suggests that a modest subset of these proteins (~1/4-1/3) are derived from the accessory glands and ejaculatory bulb. Reassuringly, the majority of these carry a secretory peptide signal. However we also found that many proteins are more ubiquitously expressed, with nearly half showing highest expression in the gonadectomized carcass. We speculate that a sizable number of identified ejaculate proteins are not playing direct reproductive roles but rather “tag along” with *bona fide* ejaculate proteins as part of the secretion process. Future efforts to characterize ejaculate proteins in this species group will reveal the functional significance of these “non-reproductive” proteins for reproductive interactions.

Finally, we assessed the utility of different species query databases for protein identification and quantitation using data dependent acquisition comparing the proteomes of closely related species. Abundance estimates varied considerably between databases and showed significantly lower correlations when using alternate species query databases (Fig. 5b), suggesting alternate species query databases provide less accurate protein abundance estimates. Amino acid substitutions may also result in less accurate protein identification and quantitation as mass-spectra are not assigned appropriately to orthologous proteins. Indeed, we found that proteins with faster rates of molecular evolution were more likely to be estimated at different abundances when using alternative databases (Fig. 5c). Furthermore, proteins without reciprocal orthology were found at lower abundance (Fig. 5d). This result shows that the detection bias against low abundance proteins in DDA may be compounded when using alternative species databases with no close ortholog to the species of interest.

Despite only retaining ~65%. of all proteins for comparison, our combined database approach was sensitive for detecting differential abundance between species. We identified similar, if not greater, proportions and absolute numbers of differentially abundant proteins between species when using the combined database. Our results show that compiling species-specific abundance data proves a useful and sensitive approach for cross-species comparisons.

## Abbreviations

DDA: data dependent acquisition
FDR: false discovery rate
FRT: female reproductive tract
GO: gene ontology
PMR: postmating female response
QTL: quantitative trait locus/loci
SFP: seminal fluid proteins
TMT: tandem mass tag

## O. Data availability

All code and analysis are available on [GitHub](https://martingarlovsky.github.io/virilisProteomics/). Supplemental data are available on DRYAD. Proteomic data have been deposited to the [ProteomeXchangeConsortium](http://proteomecentral.proteomexchange.org/) via the PRIDE partner repository (85) with the identified XXXXXXXX.

## P. Author contributions

MDG and YHAB conceived the study, collected the data, performed analyses and wrote the manuscript.

## ACKNOWLEDGEMENTS

We would like to thank Mike Deery, Yagnesh Umrania and Renata Feret from the Cambridge Proteomics facility for technical assistance, and James L. Hougland and Michelle A. Sieburg for use of a spectrophotometer. Eric Sedore and Larne Pekowsky from the Syracuse University HTC Campus Grid and NSF award ACI-1341006 provided computing services. Members of the Center for Reproductive Evolution provided helpful discussion and feedback about the project. This work was funded by funds from Syracuse University to YHAB.

## Supplementary Note 1: Supplementary information

**Fig. S1.**
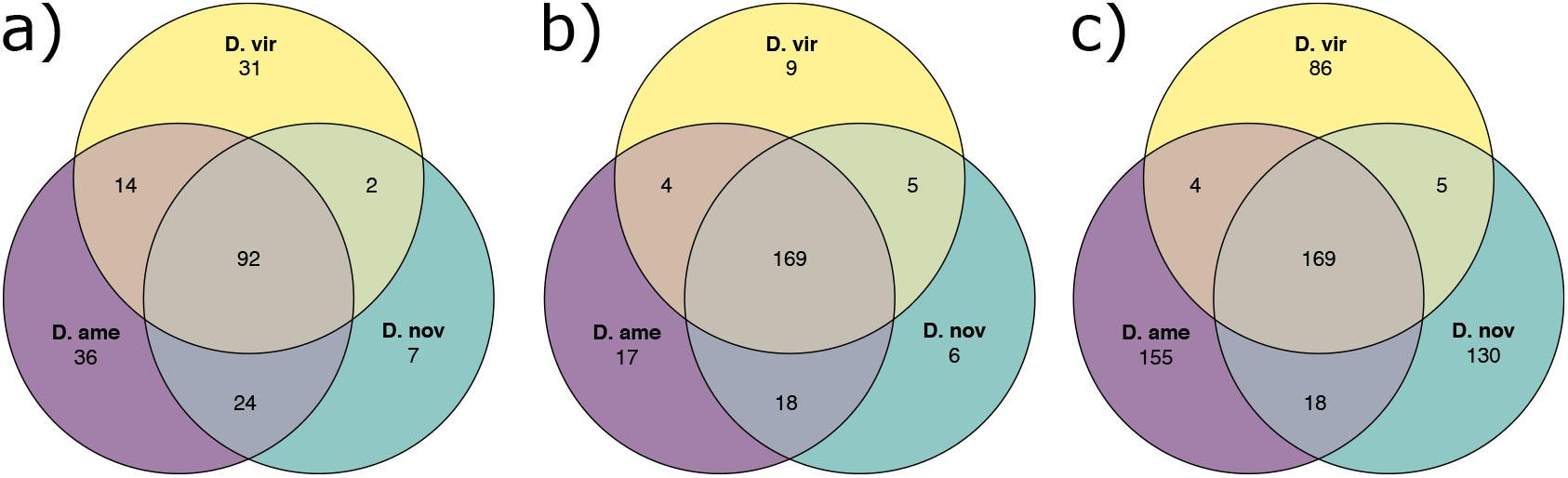
a) Shared and unique ejaculate candidates between species using the combined database. b) Shared and unique ejaculate candidates with orthologs between species using each species database. c) Shared and unique ejaculate candidates between species, including those without orthologs, using each species database.

**Fig. S2.**
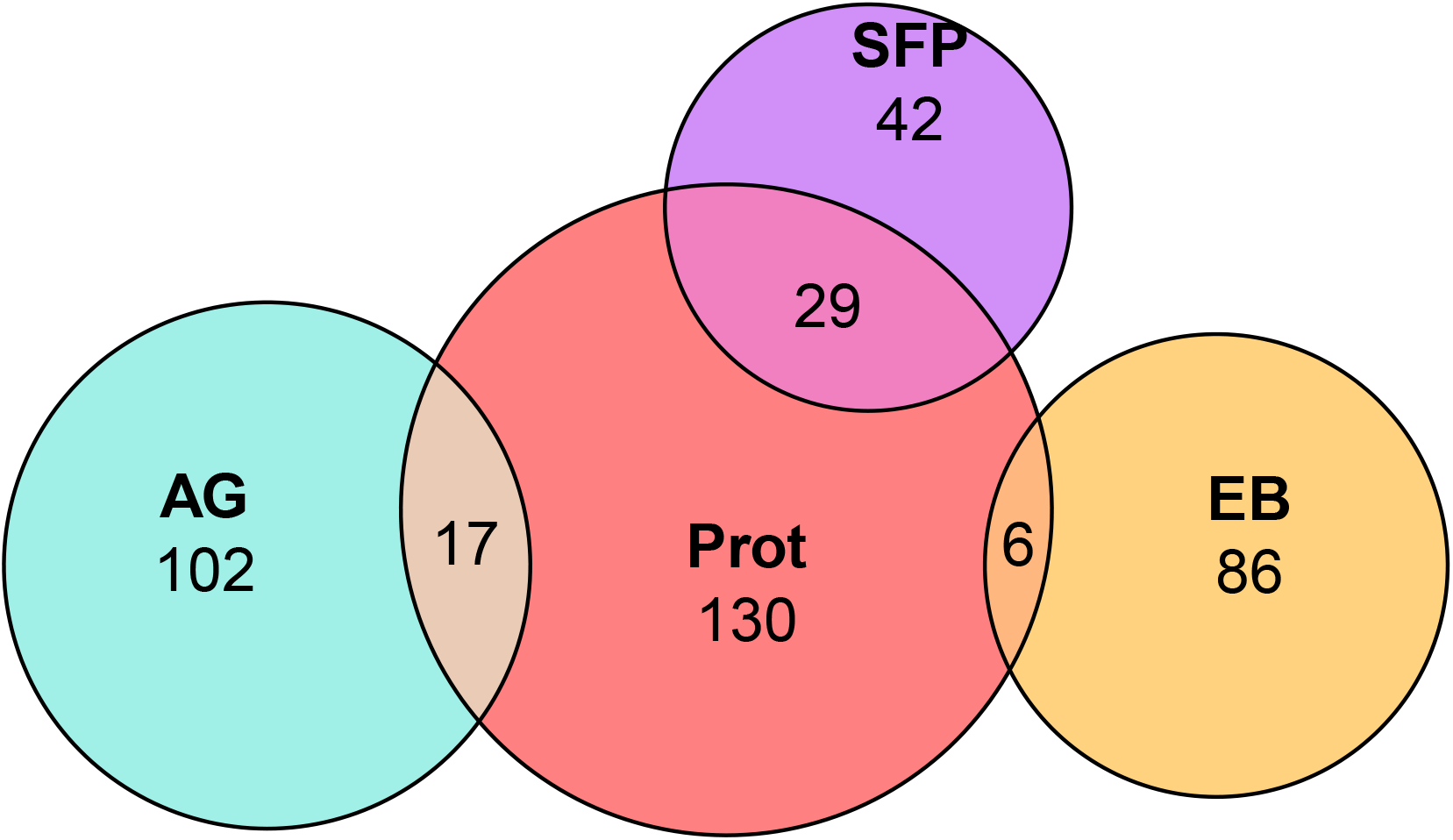
Overlap between proteins identified in the current study (Prot) with accessory gland (AG), ejaculatory bulb (EB), or SFPs identified by Ahmed-Braimah et al. (42).

**Fig. S3.**
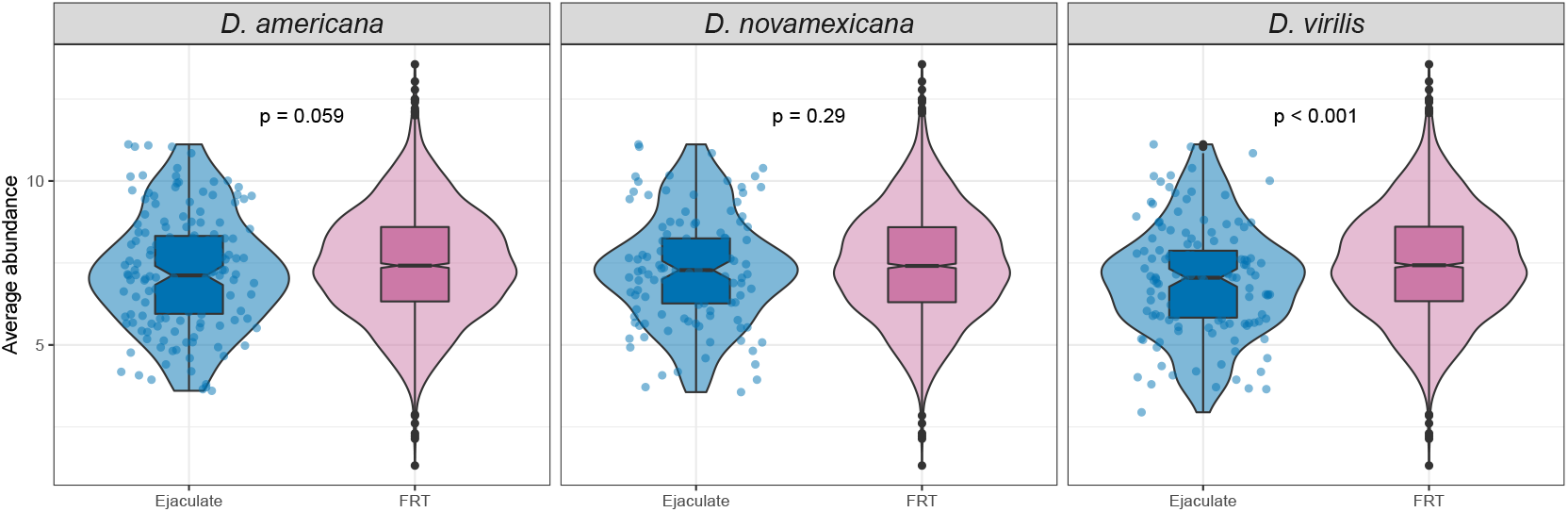
Abundance of ejaculate candidates vs. remaining female reproductive tract (FRT) proteins in each species. Points are individual proteins. P-values obtained from Mann-Whitney U tests.

**Fig. S4.**
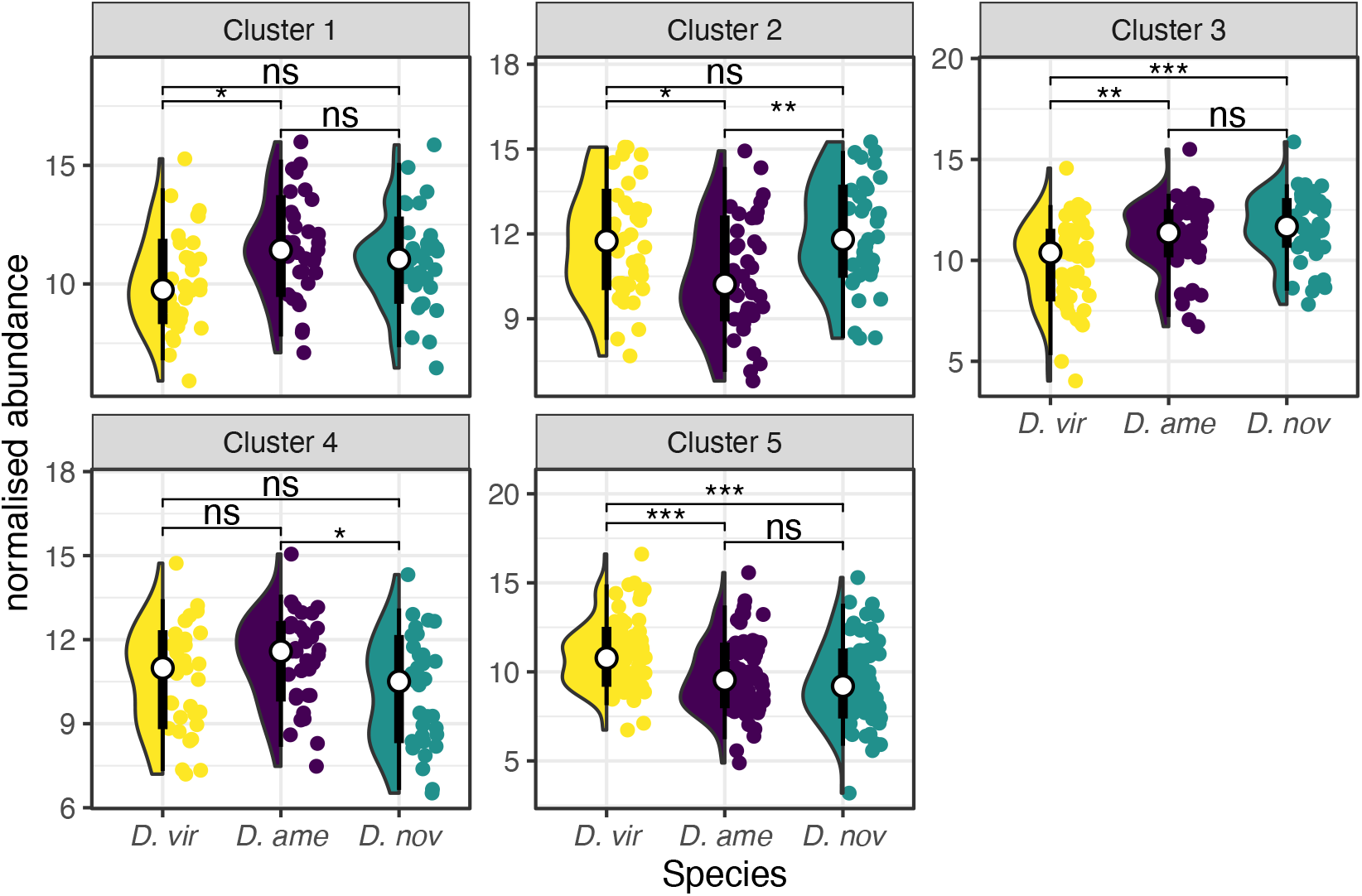
Abundance of ejaculate candidates in each cluster. K-means clustering was used to separate proteins in to k = 5 clusters, which distinguished groups of proteins differing in abundance between species. White points show the mean with thick and thin black bars showing the 66% and 95% confidence intervals, respectively. *P*-values represent results from post-hoc Tukey’s honest significant difference tests after analysis of variance performed on each cluster; ns: non-significant; *: *p* < 0.05; **: *p* < 0.01; ***: *p* < 0.001.

**Fig. S5.**
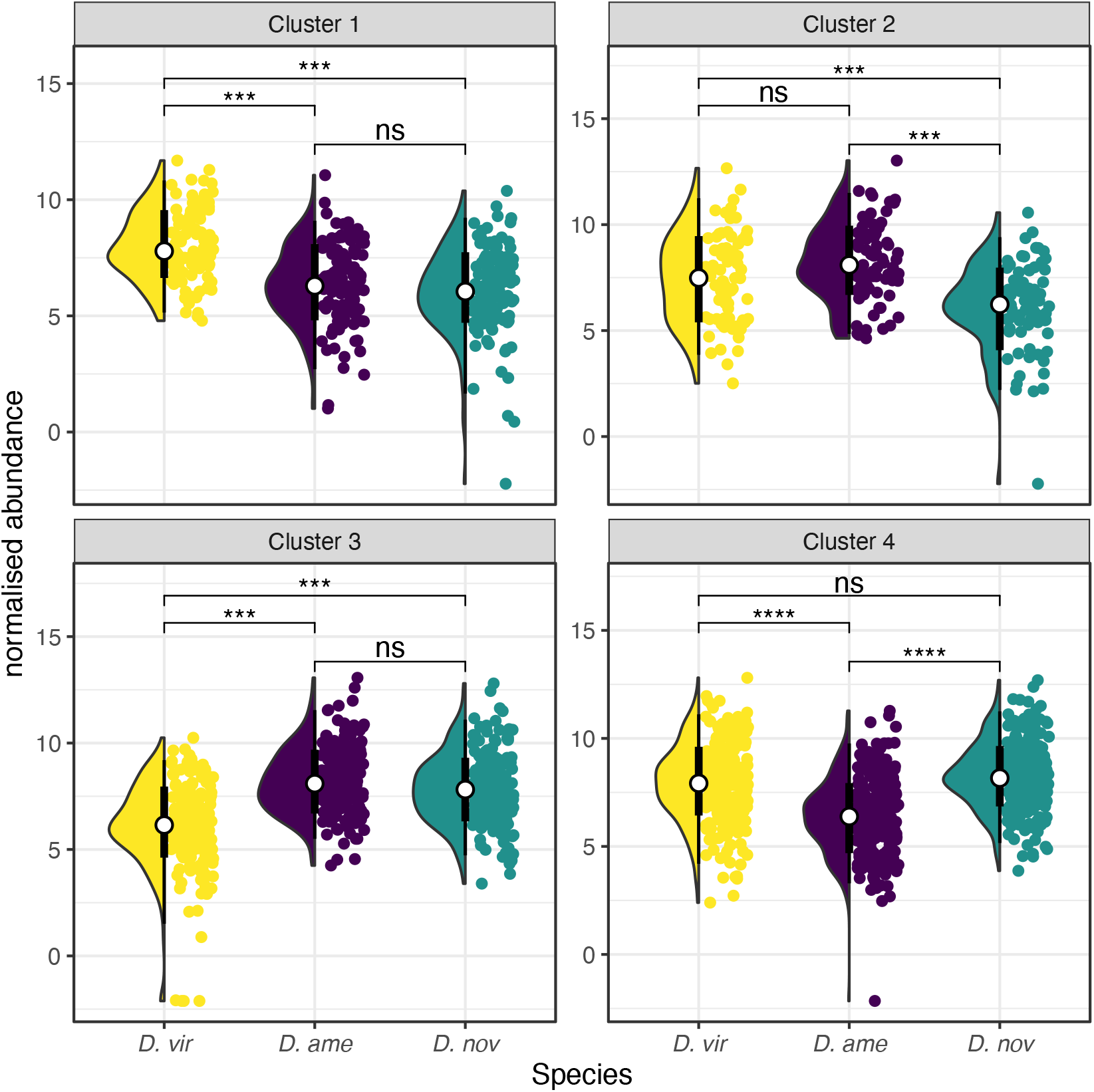
Abundance of female reproductive tract proteins in each cluster. K-means clustering was used to separate proteins in to k = 4 clusters, which distinguished groups of proteins differing in abundance between species. White points show the mean with thick and thin black bars showing the 66% and 95% confidence intervals, respectively. *P*-values represent results from post-hoc Tukey’s honest significant difference tests after analysis of variance performed on each cluster; ns: non-significant; *: *p* < 0.05; **: *p* < 0.01; ***: *p* < 0.001.

**Fig. S6.**
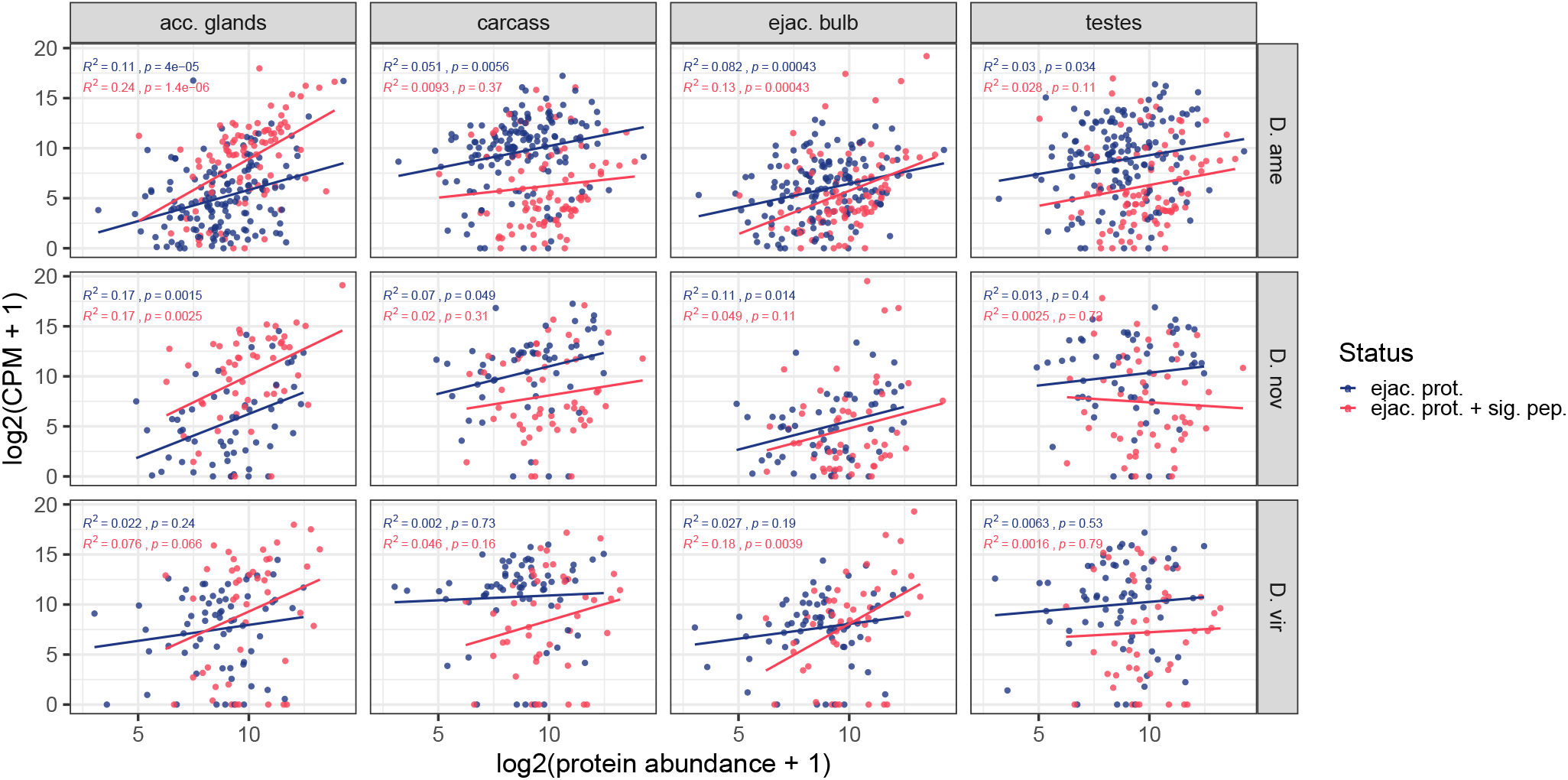
Correlation between mRNA abundance and protein abundance for ejaculate proteins across the three species and four male tissues: accessory glands, ejaculatory bulb, testes, and gonadectomized carcass. Correlation coefficients and *p*-values are indicated within each panel and separately for ejaculate proteins and ejaculate proteins with a predicted signal peptide

**Fig. S7.**
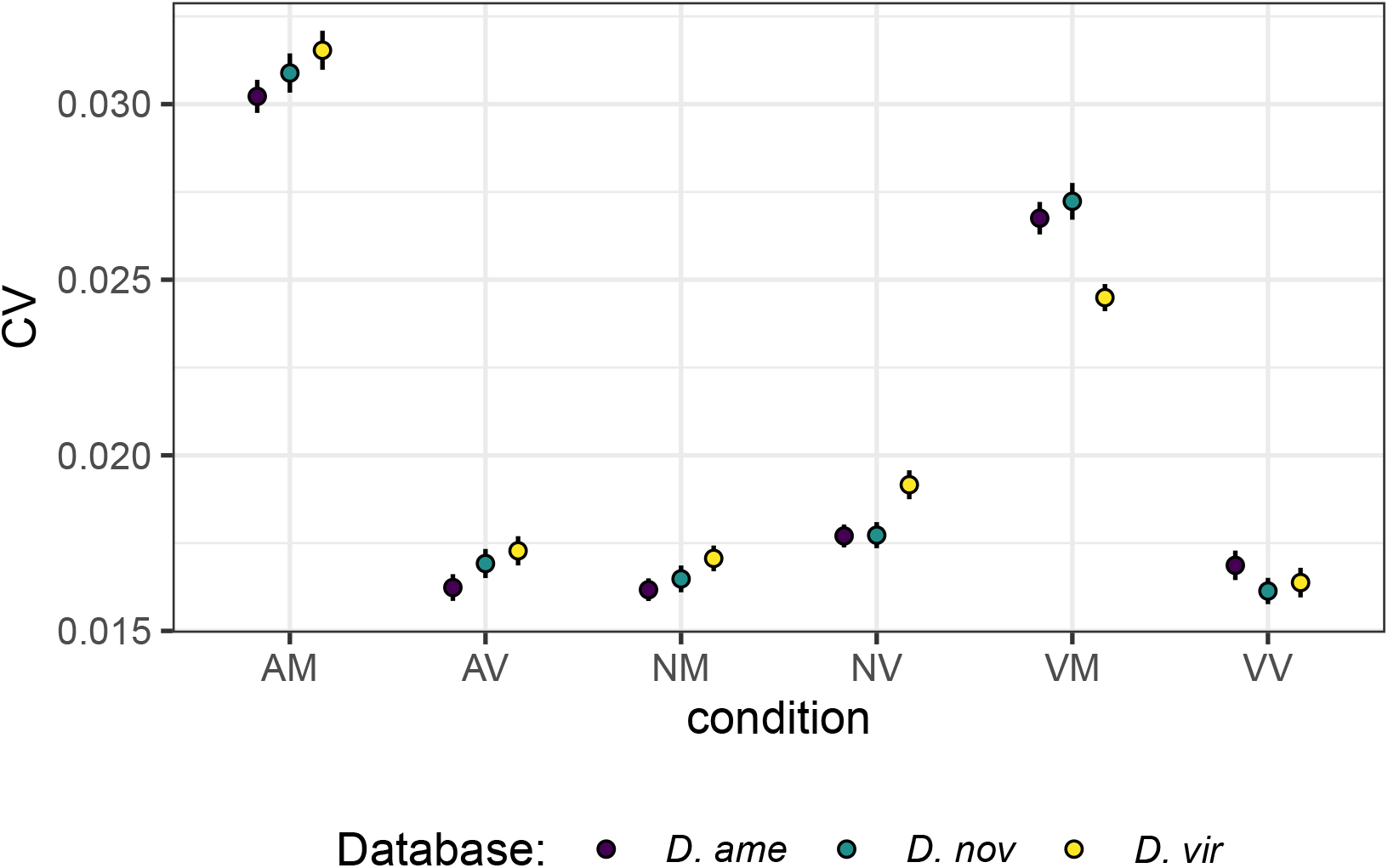
Coefficients of variation (CV) calculated for each treatment (mated vs. virgin for each species) using each species database.

**Fig. S8.**
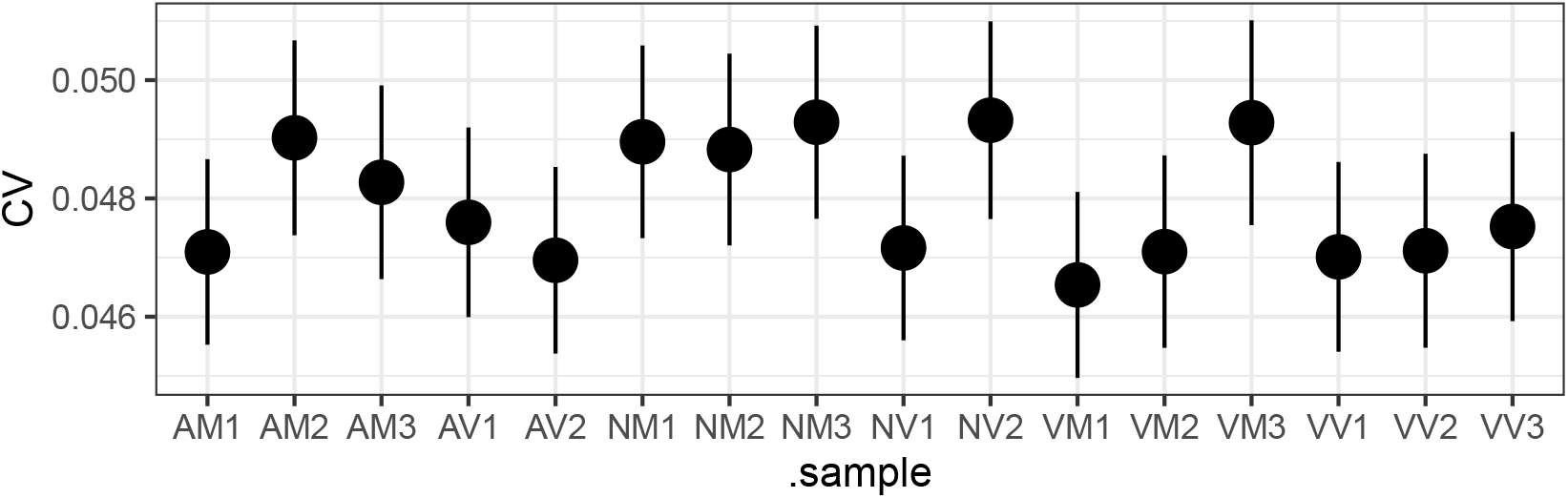
Coefficients of variation (CV) calculated for each replicate across databases.

## Notes

https://martingarlovsky.github.io/virilisProteomics/

### Competing Interest Statement

The authors have declared no competing interest.

